# Systematic comparison of risky choices in humans and monkeys

**DOI:** 10.1101/2023.02.07.527517

**Authors:** Leo Chi U Seak, Simone Ferrari-Toniolo, Ritesh Jain, Kirby Nielsen, Wolfram Schultz

**Affiliations:** Department of Physiology, Development and Neuroscience, University of Cambridge, Cambridge CB2 3DY, United Kingdom; Management School, University of Liverpool, Liverpool L697ZY, United Kingdom; Division of the Humanities and Social Sciences, California Institute of Technology, Pasadena CA 91125, USA

**Keywords:** Independence axiom, utility, risk, choice

## Abstract

The past decades have seen tremendous progress in fundamental studies on economic choice in humans. However, elucidation of the underlying neuronal processes requires invasive neurophysiological studies that are met with difficulties in humans. Monkeys as evolutionary closest relatives offer a solution. The animals display sophisticated and well-controllable behavior that allows to implement key constructs of proven economic choice theories. However, the similarity of economic choice between the two species has never been systematically investigated. We investigated compliance with the independence axiom (IA) of expected utility theory as one of the most demanding choice tests and compared IA violations between humans and monkeys. Using generalized linear modeling and cumulative prospect theory (CPT), we found that humans and monkeys made comparable risky choices, although their subjective values (utilities) differed. These results suggest similar fundamental choice mechanism across these primate species and encourage to study their underlying neurophysiological mechanisms.

## Introduction

Risky decision making has been investigated for more than 200 years (Stigler, 1950) and remains a popular research topic (Blavatskyy et al., 2022; Bujold et al., 2022; Frey et al., 2017; Ruggeri et al., 2020; Yang et al., 2022). While economists showed interest in building mathematical models to explain risky decisions and relating them to society (Moscati, 2016; Ruggeri et al., 2020; Schneider & Day, 2018), neuroscientists investigated the neural mechanism of decision making. To bridge the gap between the two directions of research, neuroeconomic studies by neuroscientists and economists have shown insightful results regarding the neural basis of risky decision-making (Bossaerts & Murawski, 2015; Konovalov & Krajbich, 2016; Serra, 2021).

In neuroeconomic research, the model organisms used by economists and neuroscientists are usually different. Economists tend to focus on human behavior and therefore conduct almost all studies with human participants (Addessi & Bourgeois-Gironde, 2020). Human neuroeconomic studies commonly use neuroimaging. While a powerful tool for studying reward value and economic decision making, the temporal and spatial resolution limits the information one can obtain from the experiments. On the other hand, more precise single-neuron studies using electrophysiology or calcium imaging are restricted to specific human patients (Nourski & Howard, 2015). Therefore, experiments on closely related species are important for studying subjective reward value and economic decision making.

While many animal species are suitable for neuroeconomic studies, neuroscientists often use rhesus macaque monkeys because the animals can understand complicated tasks and are phylogenetically closely related to humans (Brosnan, 2021; de Petrillo & Rosati, 2021). Recently, many important neuroeconomic discoveries were made in monkeys. For example, neurons in monkey orbitofrontal cortex (OFC) encode type, magnitude, probability and subjective value of reward (Ballesta et al., 2020; Padoa-Schioppa & Assad, 2006; Pastor-Bernier et al., 2019; Tremblay & Schultz, 1999), amygdala neurons encode emotional and social choices (Grabenhorst et al., 2019), and dopamine neurons encode utility and update value (Lak et al., 2014; Stauffer et al., 2014). These important studies help us to understand the neuronal mechanism of value and choice. While both neuroscientists and economists conducted valuable studies on risky decision-making, substantial gaps remain because of their failure to directly compare risky decision making between humans and monkeys. Without this information, researchers would not know whether humans and monkeys perform risky choices in a similar or different way.

Comparisons of decision making between humans and monkeys are largely limited to literature reviews (Addessi & Bourgeois-Gironde, 2020; Bourgeois-Gironde et al., 2021), but direct experimental comparisons have not been performed. In monkeys, studies demonstrated that monkeys maximize expected utility (Ferrari-Toniolo et al., 2019; Stauffer et al., 2015), cooperate with others (Grabenhorst et al., 2019), show loss aversion (Chen et al., 2006), and exhibit different risk attitude under different conditions (Ferrari-Toniolo et al., 2019; Pelé et al., 2014). In humans, on the other hand, studies investigated mathematical models and the influence of culture, education and social norms on risky decision making (Blavatskyy et al., 2022; Nielsen & Rehbeck, 2022; Ruggeri et al., 2020). By contrast, only a few behavioral studies that included both humans and non-human primates focused on simple social and economic games (Brosnan et al., 2011, 2012, 2017; Duguid et al., 2014; Farashahi et al., 2019; Möller et al., 2022), but not on quantifiable risky choices that are most commonly investigated in neuroeconomic research. Moreover, these studies tested only small choice sets, which limits the generality of the comparison between species.

Here, we investigated the similarity of risky choices between humans and monkeys, testing the independence axiom (IA) of expected utility theory with exactly the same design and settings between the two species. The IA states that extending both choice options of a binary option set by a common outcome should not change the participant’s preference. However, our previous work had shown violations of the IA separately in humans and monkeys (Ferrari-Toniolo et al., 2022; Jain & Nielsen, 2020). Therefore, by using this axiom as a test, we systematically compared risky choices between the two species.

## Methods

### Animals

Two rhesus macaques (Macaca Mulatta) weighing 12.65 kg and 13 kg were used in this experiment. The monkeys were born in captivity at the UK Medical Research Council’s (MRC) Centre for Macaques (CFM). Monkey A (“Aragorn”) and Monkey T (“Tigger”) were pre-trained with visual stimuli and similar joystick tasks before the experiment. The protocol was approved by the Home Office of the UK and the experiments were continuously regulated by institutional (University of Cambridge) and national officers including the UK Home Office Inspector, the University of Cambridge Biomedical Services (UBS) Named Veterinary Surgeon (NVS), and the UBS Named Animal Care and Welfare Officer (NACWO). Monkey T was implanted with a recording chamber and a headpost before the experiment for other neuronal recording tasks.

### Experimental design for monkeys

During the experiment, each monkey sat in a chair (Crist instruments) and chose between two gamble options using a cursor driven by a left-right joystick. The two gamble options were presented on a computer monitor 50 cm in front of the animal. In each option, reward magnitude (varying between 0 and 0.5 ml water) and probability (varying between 0 and 1) were represented by the height and width of a horizontal bar, respectively (Fig. 1A). At the beginning of each trial, a white fixation cross appeared at the center of the monitor, and the cursor was displayed to facilitate centering the left-right joystick. After two gamble stimuli were shown, the animal chose using the joystick. Further details of the experiment can be found in our previous studies using similar setups (Ferrari-Toniolo et al., 2019, 2022).

**Figure 1.**
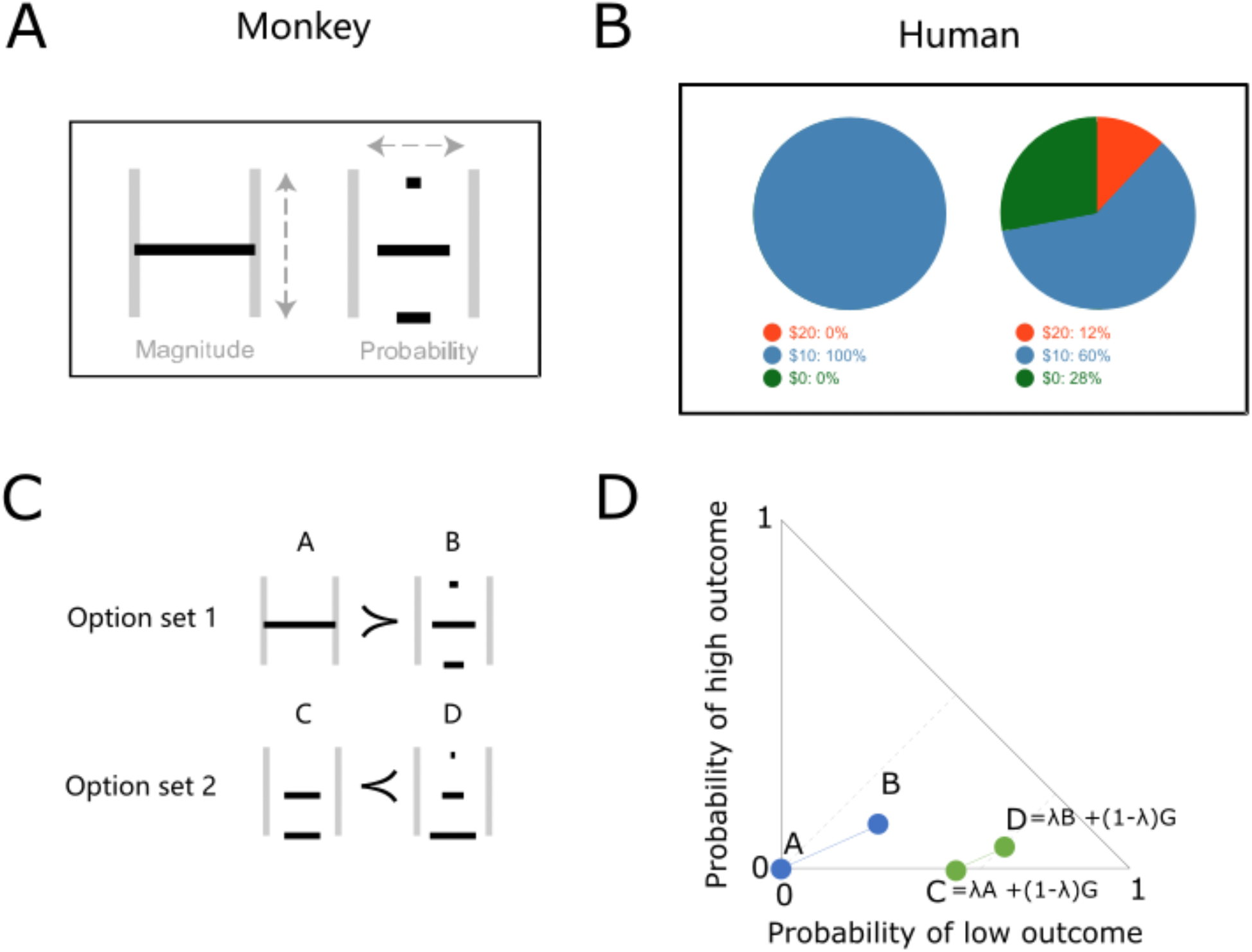
Experimental design. (A) Example stimuli shown on the computer monitor to monkeys. Width and height of horizontal bars represent outcome probability and magnitude, respectively. The example shows options A and B. (B) Example stimuli shown to human participants (options A and B). (C) Stimuli showing two option sets for independence axiom testing. Option A had only the middle outcome with p = 1. Option C had a middle outcome with a specific probability and a low outcome of m = 0 ml and p (low outcome) = 1 – p (middle outcome). Options B and D each had three outcomes with specific probabilities. Preferring option A (A > B) and option D (C < D) represents an example violation of the independence axiom. (D) Marschak-Machina triangle with the two option sets shown in panel C. Dot positions in the triangle represent the outcome probabilities for options A, B, C and D. The probability of the middle outcome is 1 - p (low outcome) – p (high outcome). Thus, option A is a safe option with p = 1.0 of getting the middle outcome.

We set up our list of tests according to the IA:

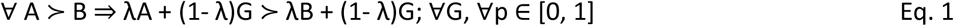

where A and B represent the two original options; λ (lambda) represents the probability of the original option in the new gamble option; G represents a gamble that commonly extends both original choice options. In this study, option A was always a safe option with a middle reward magnitude (0.25 ml); option B was a three-outcome (0, 0.25ml, 0.5ml) option. Option A and B were extended by a new gamble G according to Eq.1, thus forming options C and D: option C = λA + (1-λ)G, and option D = λB + (1-λ)G. For gamble G, we set reward magnitude to 0 (no reward) and used three different probabilities λ: 0.75, 0.5 and 0.25, together with the original test set (λ = 1), thus forming four different test settings. All choice options were pseudorandomly intermixed.

### Experimental design for humans

We conducted 180 different online tests on 126 human participants. Each participant performed 34.254 ± 7.2354 tests for each of the 4 lambdas (mean ± Standard Deviation). These tests were selected from a previous study (Jain & Nielsen, 2020) and had similar probability distributions as used for monkeys, including option A having only one reward amount delivered with p = 1.0 (Fig. 1B). Specifically, the monkeys were tested with reward magnitudes of 0 ml, 0.25 ml and 0.5 ml, and the human participants were tested with reward magnitudes of $0, $10 and $20. All tests were pseudorandomly ordered with each participant. In each trial, two different options were shown to the participant on a computer monitor. Each option was shown as a pie chart, indicating the probability of reward, together with numbers indicating each option. Participants were paid for one randomly selected choice after completing the session, as described in previous economic studies (Azrieli et al., 2020).

### Statistical test of IA violations

To test IA violations, we assessed the probabilities P(AD) and P(BC) that represent the two directions of violations across sessions in each monkey and across individual human participants. The violation AD indicates an Allais-type reversal: a participant who prefers the safe option A in option set AB prefers the “riskier” option D in option set CD, which can be stated as: p (A|{A,B}) > 0.5 and p (D|{C,D}) > 0.5. By contrast, the violation BC indicates a reverse Allais-type reversal: a participant who prefers option B in option set AB prefers option C in option set CD, which can be stated as: p (B|{A,B}) > 0.5 and p (C|{C,D}) > 0.5. The probability of a preference reversal in each direction, P(AD) or P(BC), was computed as proportion of monkey choice sessions or as proportion of single-shot human choices that switched in the corresponding direction.

To confirm the preference changes across sessions in each monkey and across individual human participants, we performed two binomial tests for the two option sets (AB, CD). Significant preference changes in both option sets would be counted as a (strong) violation of IA (P < 0.05; one-sided binomial test).

### Comparison of risky choices between humans and monkeys using a General Linear Model (GLM)

We used Matlab to build GLMs (fitglm with normal distribution and linear link function) with human data to predict monkey behavior. The GLMs predicted the probability y of choosing option B or D based on three regressors, namely lambda (λ in Eq. 1), the probability p of obtaining the highest outcome, and the probability p of obtaining the lowest outcome) (fig. 1C):

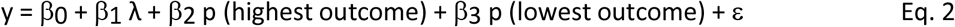

After fitting the model (Eq. 2) to the human data, we entered the reward probabilities for the monkeys into Eq. 2 and predicted the probability of choosing option B or D. Inversely, after fitting the model to the monkey data, we entered the reward probabilities for the humans into Eq. 2 and predicted the probability of choosing option B or D. The predicted probabilities of choices were compared with the actual choice probabilities using Pearson’s correlation analysis. We compared the predictions of the human-fitted model to each monkey separately, and we compared the prediction of the monkey-fitted model to all humans’ data. To further investigate how each variable contributed to the GLM, we also assessed the beta coefficients (slope) for the regressors of the models. In order to fit the single-shot human choices, we calculated the GLM using the binomial distribution and logit link function, repeating the procedure for each human participant and for each monkey session. We then plotted the betas of each model in a three-dimensional scatter plot after normalization (deducting the minimum value and log10 transformation) and after removing outliers with more than three median absolute deviations (Fig. 3).

**Figure 2.**
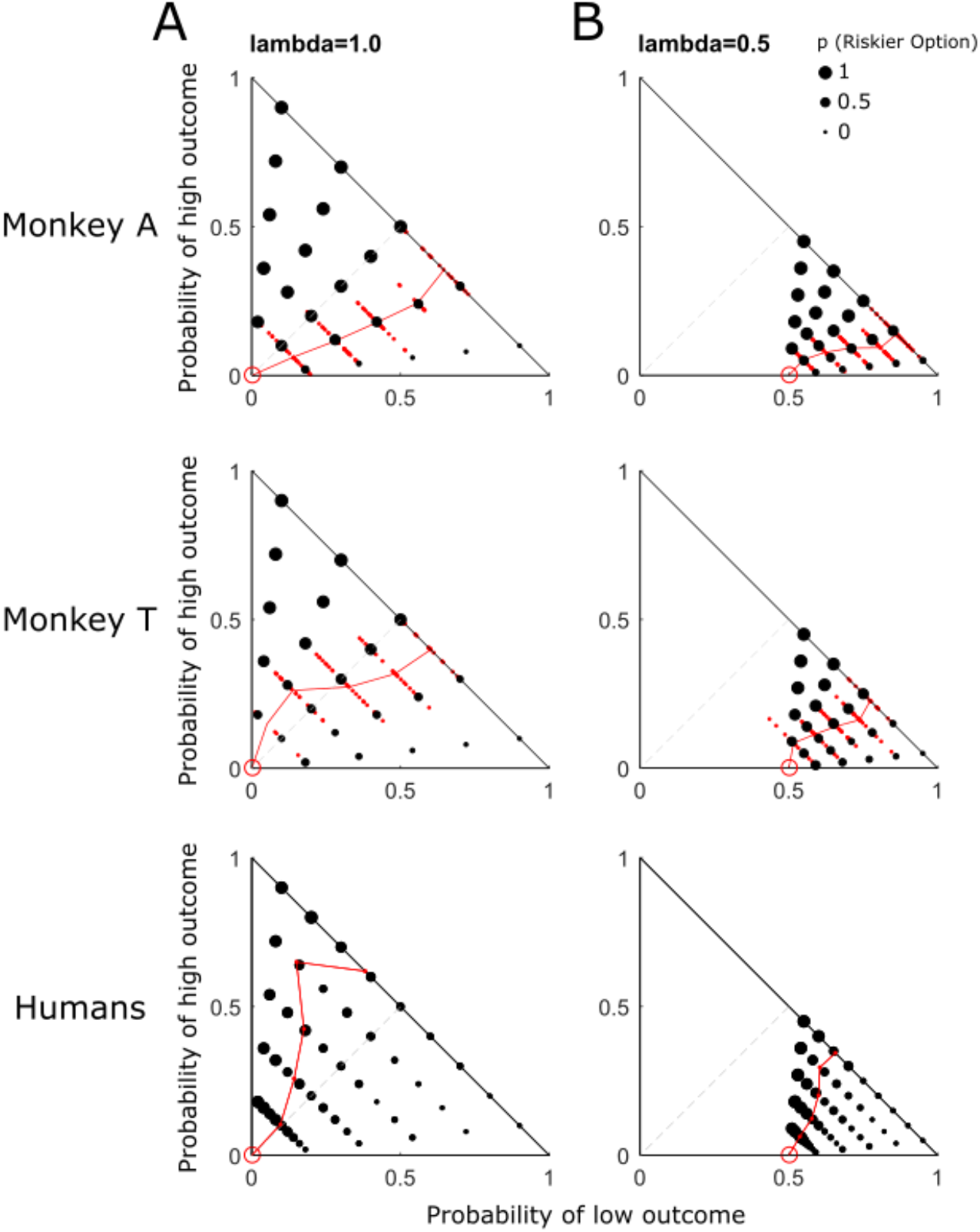
Differential risk attitude across reward probabilities in the Marschak-Machina triangle. (A) Choice between options A and B (λ = 1.0). (B) Choice between options C and D (λ = 0.5).

**Figure 3.**
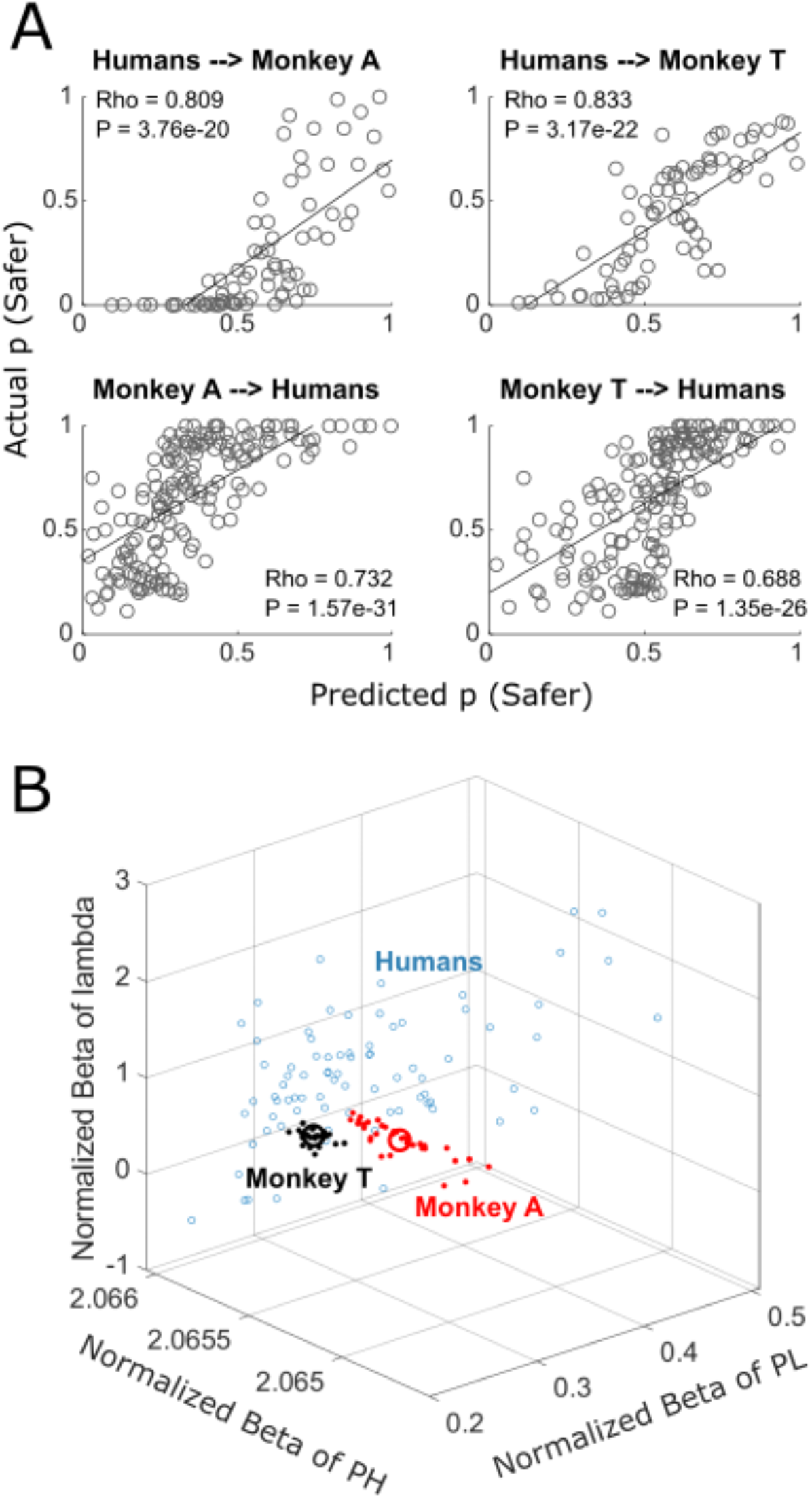
General Linear Models (GLM’s) of human data (probability of safer choices) predict monkeys’ behavior and vice versa. (A) Correlations of choice probabilities predicted by GLM and actual choices. Each circle represents one test in the Marschak-Machina triangle (see Figs. 2 and S1). Pearson’s Rho, least squares lines. (B) Beta coefficients of GLMs fitted to data from humans (each participant; blue open circles) and monkeys (each session; red or black dots for Monkey A and Monkey T, respectively). The average betas for monkeys are represented by large open circles (red for Monkey A and black for Monkey T). PH and PL represent reward probability: PH represents p (high outcome) and PL represents p (low outcome) for the riskier option.

### Cumulative prospect theory model

Similar to previous studies (Ferrari-Toniolo et al., 2019, 2022), we used a softmax function to describe the choices as follows: the probability (p) of choosing an option M over an option N, given the option set MN, was defined as:

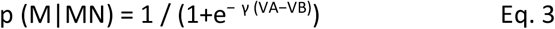

where γ is the noise parameter and V is the prospect value (i.e. the subjective value as defined in the cumulative prospect theory model. We defined the prospect value (V) using the utility function (u) and probability weighting function (w) in a cumulative form (Kahneman & Tversky, 1979; Tversky & Kahneman, 1992):

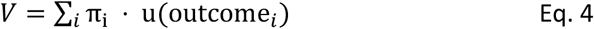

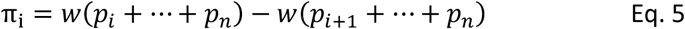

where n is the number of outcomes and i is the corresponding current outcome (ordered from worst to best).

As in previous studies (Ferrari-Toniolo et al., 2019; Hsu et al., 2009), we used a power function as utility function, with parameter ρ (>1 convex, <1 concave):

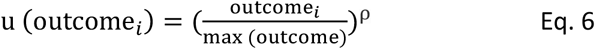

For the probability-weighting function, we used the two-parameter Prelec function with (α, β):

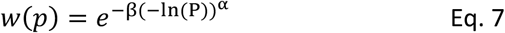

The α and β represent the shape (α < 1: inverse S shape, α > 1 regular S shape) of the probability-weighting function and the position of the inflection point, respectively.

Similar to our previous work (Ferrari-Toniolo et al. 2022), we estimated the parameters of the utility function and weighting function by maximum likelihood estimation (MLE). Choice data from each human participant or each monkey session were entered into Matlab (fminsearch function) to estimate the four parameters (γ, ρ, α, β).

## Results

### Experimental design

We used a common experimental design to compare risky decisions between humans and monkeys. For monkeys, we showed two stimuli on a computer monitor at 0.5 - 1.0 s after appearance of a central fixation cross. Subsequently, the animal chose between the two options using a joystick and received the reward 1s later. As shown in Fig. 1A, each of the option stimuli contained one, two or three horizontal bars whose width and height represented reward probability (p = 0 - 1) and magnitude (m = 0 - 0.5 ml), respectively. For example, a full-width horizontal bar in the middle would represent a safe option (p = 1) of a middle reward (m = 0.25 ml) (Fig. 1A right). For humans, we presented each option as a pie chart and numbers that indicated reward amount (US$ 0 - 20) and probability (p = 0 - 1) (Fig. 1B).

Each test of the independence axiom (IA) employed two option sets. As shown in the example stimuli (Fig. 1C) and Marschak-Machina triangle (Fig. 1D), one option set (options C and D) was derived from the original option set (options A and B) according to the definition of the IA (Eq. 1). To comply with the IA, the participant’s preference should not change (e.g. if option A is preferred to option B, then option C should also be preferred to option D). In order to test the IA fully, we varied the options widely across the full Marschak-Machina triangle in both humans and monkeys.

### Characteristics of risky choice in humans and monkeys

The monkeys performed a total of 92 tests in each daily session (23 tests for each of the 4 lambdas) (30,997 trials in 34 sessions for Monkey A; 11,492 trials in 26 sessions for Monkey T). On average, each session consisted of a total of 911.68 ± 178.69 trials in Monkey A, and 442.00 ± 123.93 trials in Monkey T (mean ± Standard Deviation).

Both monkeys preferred options A and C more (compared to the respective options B and D) when the probability of the high outcome of options B and D decreased, as shown by the decrease of dot size towards the bottom in the Marschak-Machina triangle (Fig. 2A, B). This observation suggested that the animals understood the stimuli and the task.

The size of black dots in the Marschak-Machina triangles represents the probability of choice in 34 daily sessions for Monkey A, 26 daily sessions for Monkey T, and 126 human participants. Red solid dots show choice indifference points (IPs); red lines show indifference curves from averaged session IPs in monkeys, and averaged IPs of all human participants.

Using the softmax function (Eq. 3), we estimated choice indifference points between option A (P (middle outcome) = 1.0; red circles in Fig. 2A) and option B (three outcomes; black dots) in each monkey in one daily session. The three lambda values tested (λ = 0.25, λ = 0.5, λ = 0.75) allowed three comparisons against λ = 1.0 (option set AB in Fig. 1C). Then we applied the three lambda values (0.25, 0.5 or 0.75) to options A and B to obtain option C (p (middle outcome) = 0.25, 0.5 or 0.75; p (low outcome) = 1 - p (middle outcome)) and option D (three outcomes) (Fig. 1C, D) and estimated choice indifference points between option C (red circles in Fig. 2B) and option D in each monkey. All indifference points are shown as small red dots in Figs. 2 and S1.

Interestingly, the probability of risky choices and the shape of indifference points differed between the two animals (Fig. 2A, B, top vs. middle). The difference was also apparent in the indifference curves (IC) connecting the averaged indifference points across sessions (red lines). Monkey A showed convex ICs with λ = 1.0 but inverse-S-shaped ICs with λ = 0.5. Monkey T showed inverse-S-shaped ICs with both λ =1.0 and λ =0.5. These data demonstrated subject-specific risk attitudes.

Many economics tests on humans use only one trial per participant in a larger number of participants (as compared to many repeated trials in much fewer individual monkeys). Accordingly, we performed 180 tests in our 126 human participants (45 tests for each of the 4 lambdas without any repetition of trial. As shown in Fig. 2, our human participants showed a similar trend (choosing the less risky option with higher probability of getting the high outcome), although their ICs differed substantially from those of our monkeys.

### GLMs reveal similarity of risky choices across species

To quantitatively measure the similarity of risky choices between humans and monkeys, we first fitted a generalized linear model (GLM; Eq. 2) to the data of one species and then used the model to predict the other species’ behavior. Fig. 3A shows strong significant correlations between the predicted probability of choices (choosing safer option; same option set across sessions in monkey, across participants in human) and the actual probability of choices. The GLM fitted to the human data top panel) predicted well the behavior of both monkeys (Rho = 0.809, p = 3.76 × 10^-20^ for Monkey A; Rho = 0.833, p = 3.17 × 10^-22^ for Monkey T). Vice versa, the GLM fitted to the data of one monkey (bottom panel) predicted well the behavior of the humans (Rho = 0.732, p = 1.57 × 10^-31^ for data by Monkey A; Rho = 0.688, p = 1.35 × 10^-26^ for data by Monkey T). These results indicated similar risky choices between the two species.

Next, we investigated whether humans’ and monkeys’ choices similarly reflected changes in the options’ parameters (probability of getting different reward magnitudes). To do so, we analyzed the beta coefficients of the GLMs. While the GLMs described the prediction of risky choices, the beta coefficients (regressors) of the GLMs indicated how much each parameter contributed to the risky choice. We fitted one GLM for each human participant and one GLM for each monkey session. As choices were not repeated in the individual human participants, we set up the model to predict single binary choices; the analysis used a binomial distribution and logit link function in the GLM. We found similar ranges of beta coefficients across the two species (Fig. 3B), suggesting similarity in risky choices between humans and monkeys.

### Choice differences between the species

We compared choice probabilities between lambda = 1 (i.e. option set AB) and other lambdas (0.75, 0.5, 0.25; option set CD) and checked for any choice shifts to identify violations. We quantified violations in two directions, the Allais paradox as originally described (Allais, 1953), and the reverse Allais paradox (Blavatskyy, 2013). Both monkeys showed significant IA violations in both directions; when preferring option A to B, they also preferred option D to C (‘Allais-type’; P < 0.05, binomial test; Fig. 4A); vice versa, when preferring option B to A, they also preferred option C to D (‘reverse Allais-type’; Fig. 4B); see Fig. S2 for additional tests in both directions. These violations differed between the two monkeys (Fig. 4, top vs. middle). Interestingly, choice variability may partly explain IA violations in monkeys (Fig. S3).

**Figure 4.**
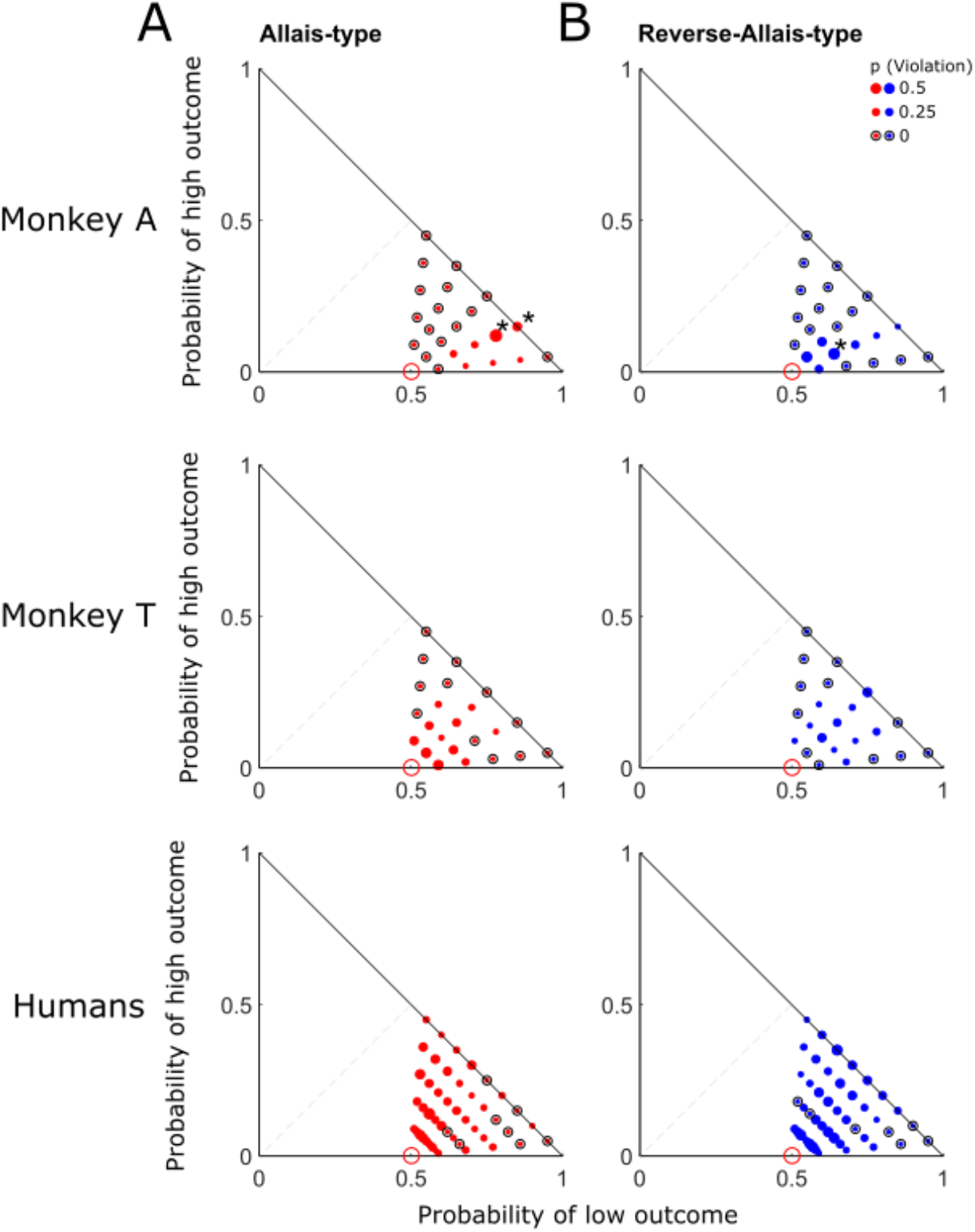
Violations of independence axiom in monkeys and humans displayed in the Marschak-Machina triangle. (A) Allais-type violations. (B) Reverse-Allais-type violations.

Dot size represents violation probability across 34 sessions in Monkey A, 26 sessions in Monkey T, and 126 human participants, respectively. Hollow black circles indicate no violation for that option set. * significance (P < 0.05; binomial test). Data shown are for lambda = 0.5; for full results, see Fig. S2.

A number of human participants violated the IA (non-null probability of violation), as shown before (Blavatskyy et al., 2022; Jain & Nielsen, 2020; Nielsen & Rehbeck, 2022), although their averaged choices failed to reveal significant IA violations (P > 0.1; binomial test) (Fig. 4 bottom). Tour surprise, the violation patterns between humans and monkeys were quite different. For example, in Fig. 4, more violations were found at the left bottom corner in the Marschak-Machina triangle in humans than in monkeys. In contrast to the GLMs fitted to the human choice probabilities (Eq. 2), the GLMs fitted to the human IA violations (Eq. 2) failed to predict our monkeys’ behavior (Rho = 0.0786, P = 0.5541, Pearson’s correlation analysis). Thus, despite the described inter-species similarities in risky choices, the more stringent IA tests nevertheless demonstrated some differences between the species.

Each black line indicates the function in each human participant or in each monkey session. The estimations used power utility functions and two-parameter Prelec probability-weighting functions (Eqs. 3 – 7). Blue lines and purple bands show averaged functions and 95% confidence intervals, respectively.

### Utility differences explain the choice differences between the species

In order to explain the differences between the two species, we fitted power utility functions and two-parameter Prelec probability-weighting functions to the choices of both humans and monkeys according to Cumulative Prospect Theory (Eqs. 3 – 7; see Methods) (Fig. 5). The two species showed very different utility functions. Specifically, the utility parameter ρ differed significantly between the two species (P = 2.0950 × 10^-13^, Wilcoxon rank sum test), whereas the shape parameter α of the probability-weighting function varied only insignificantly between the two species (P = 0.4409). The more concave utility functions in humans compared to monkeys indicated more pronounced risk avoidance in our human participants. However, some utility function in individual human participants resembled those of individual monkeys (Fig. 5, individual black lines).

**Figure 5.**
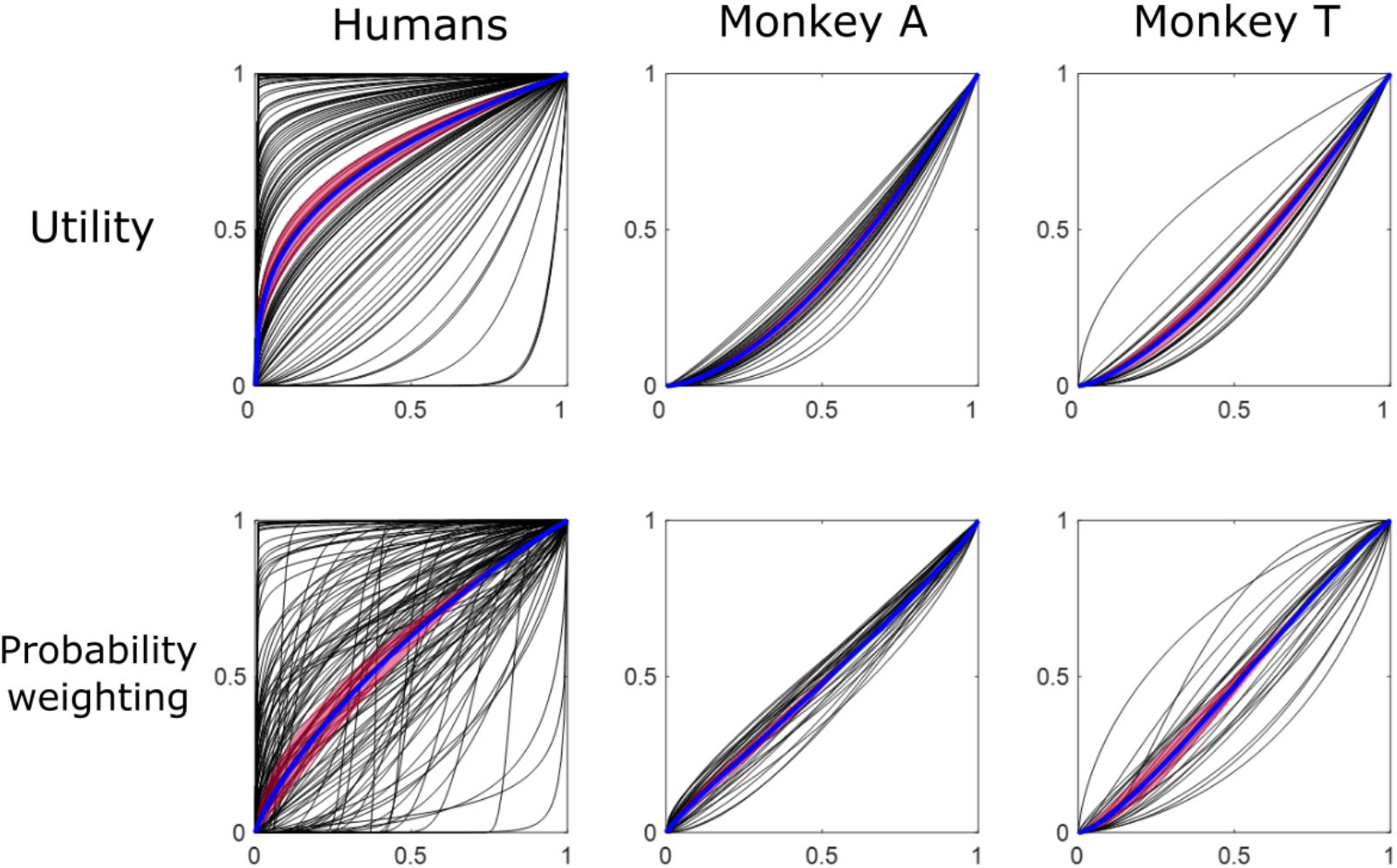
Utility and probability-weighting functions in humans and monkeys.

## Discussion

This study investigated similarities and differences in risky choices between humans and monkeys. We compared risky choices between the two species in a systematic way by using exactly the same experimental design. Although we found overall similar choices in the two species, specific tests of the independence axiom (IA) revealed substantial differences in utility functions between the two species, whereas their probability weighting functions were similar.

Current electrophysiological research in monkeys aims to elucidate neuronal mechanism of risky choices. For example, recent monkey studies demonstrated that neurons in the orbitofrontal cortex, amygdala and anterior insular cortex compute reward value in risky choices (Ballesta et al., 2020; Grabenhorst et al., 2019; Suzuki et al., 2017; Yang et al., 2022). Follow-up studies propose more advanced reinforcement learning and economic models based on neuronal recording data (Dabney et al., 2020; Serra, 2021). While these studies provided critical information about neuronal decision mechanisms, it remains unknown to which extent these results in monkeys may explain human economic decision mechanism. Surprisingly, no previous study systematically compared human choice behavior and monkey choice behavior under risk. Therefore, our results demonstrating similar risky choices in humans and monkeys may provide foundations for these neurophysiological studies.

Previous economics studies demonstrated that risk attitude depends on many factors, such as sequence of option presentation, loss or gain frames, social and cultural factors, and personality (Malenka et al., 1993; Mikels & Reed, 2009; Ruggeri et al., 2020; van den Bos et al., 2013). Even in well-controlled experimental setting, monkeys show differences in performing risk choices compared to humans, as shown in Figs. 4 and 5. Therefore, even with a strict experimental design and an absence of cultural influence, individual risk attitudes still exist that can be explained by differences in the subjective evaluation of rewards as shown by the differences of utility functions.

Our previous work showed similar violations of the IA in monkeys and humans in specific settings of the IA (Ferrari-Toniolo et al., 2022). Our current study used a far larger range of IA tests in humans and option sets that directly reflected the axiom definition in monkeys. Using this extended design, we still found violations of the IA in monkeys that occurred in both directions (Allais and reverse Allais). These results confirm that expected utility theory cannot explain all choices under uncertainty and that the IA violations are not restricted to special circumstances (common consequence and common ratio effects). The monkeys had different individual risk attitudes and violation patterns, which is consistent with previous human studies on risky decision-making (Blavatskyy et al., 2022; Ruggeri et al., 2020). It would be interesting to investigate whether monkeys’ behavior is more similar to some (but not all) humans.

In line with cumulative prospect theory, we interpreted IA violations as resulting from the subjective non-linear evaluation of reward probabilities, formally represented by the probability weighting function. On the other hand, the subjective non-linear evaluation of reward magnitudes (i.e. the utility function) cannot by itself generate IA violations but can contribute to their specific patterns. Our data revealed systematic IA violations in both humans and monkeys, suggesting a common probability weighting mechanism, while different utility functions generated different patterns of violations in the two species. Overall, these data suggest a brain mechanism for the evaluation of risky choice options that is compatible with cumulative prospect theory and, importantly, is common to humans and monkeys.

While most previous economic studies focused only on single deterministic choices (Blavatskyy et al., 2022; Blavatskyy, 2007), our study on monkeys tested choices repeatedly. The repeated choices reduced the chance of mistakes and noise that might explain some axiom violations (Blavatskyy, 2007; Hey & Orme, 1994). The probability of IA violations was positively correlated with choice variability (standard deviation of choice probability; Fig. S3), which might be explained by the observation that most violations occurred close to the indifference points and curves of the Marschak-Machine triangle (Figs. 2 and S1) (McGranaghan et al., 2022). To conclude, our extensive and inter-species study confirms the well-known systematic and subject-specific violations of the IA.

Critically, our study showed not only similarities but also differences between the two species, notably in subjective value (utility). The differences may be due to at least two factors. First, typical for monkey studies, we used large numbers of trials that provided substantial experience for the animals. By contrast, typical human studies use single-shot choices that fail to provide much experience with the tested option sets. This difference in experience between the two species may partly explain their different utility functions. Second, reward types and their amounts differed between the two species. Our human participants were tested with money, whereas our monkeys were tested with juice. Further, humans and monkeys seem to be risk seeking with small reward amounts and become gradually more risk avoiding with larger amounts, which is expressed by the curvature of utility functions that changes from convex via linear to concave (Stauffer et al., 2014; Farashahi et al., 2018). The risk-seeking with small reward amounts is recognized as the peanuts effect (Prelec, 1991). Thus, despite the difficulties of inter-species comparisons, different reward types and amounts may have contributed to the differences in utility functions. Future research may elucidate in more detail the differences between human and monkey utility functions with a more quantitative approach towards task experience and reward amount.

## Acknowledgements

We thank Aled David and Christina Thompson for animal and technical support. This study was supported by Wellcome Trust (WT 095495, WT 204811, WT 206207), European Research Council (ERC; 293549) and US National Institutes of Mental Health (NIMH) Caltech Conte Center (P50MH094258). For the purpose of Open Access, the authors have applied a CC BY public copyright licence to any Author Accepted Manuscript version arising from this submission.

## Supplementary figures

**Figure S1.**
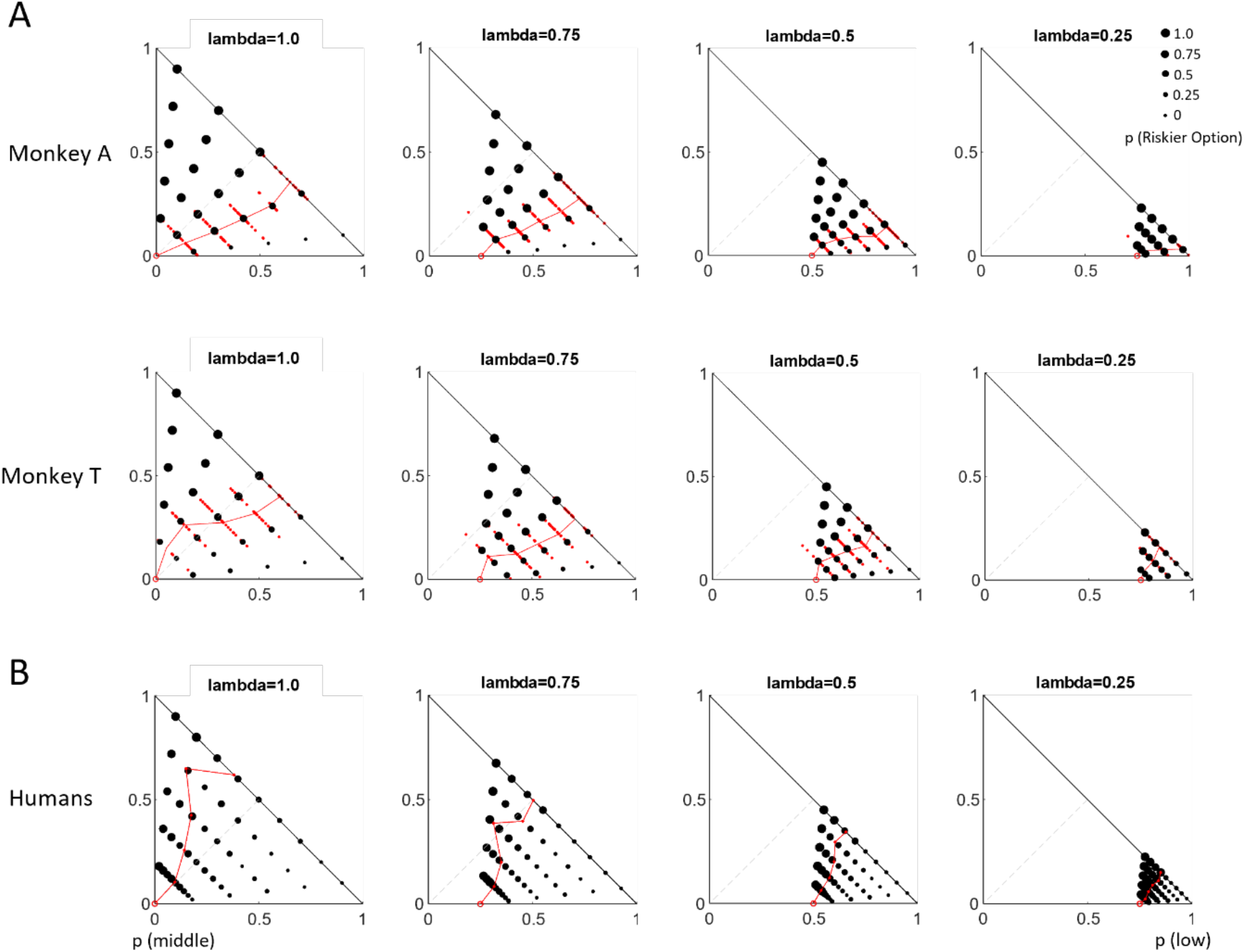
Differential risk attitude across probabilities in the Marschak-Machina triangle. (A) Choices in the two monkeys: 34 sessions for Monkey A and 26 sessions for Monkey T. (B) Choices in the 126 human participants: four safer options ($10, p = 1; $10, p = 0.75; $10, p = 0.5; $10, p = 0.25). Each panel shows the probability of choosing a risky option. Each trial offered two options, a safer option (blue solid dot) and a riskier option (one of the black solid dots). We tested four safer options (0.25ml, p=1; 0.25ml, p = 0.75; 0.25ml, p = 0.5; 0.25ml, p =0.25), as represented by the four Marschak-Machina triangles (lambda = 1.0, lambda = 0.75, lambda = 0.5, and lambda = 0.25). The probability of choosing the risky option across sessions is represented by the size of the black solid dot. Indifferent points (IP) in each session, indicating p = 0.5 of choosing the risky option (as estimated by softmax function), are represented by red dots. Red hollow circles represent average IPs across all sessions; blue dots represent the safer options.

**Figure S2.**
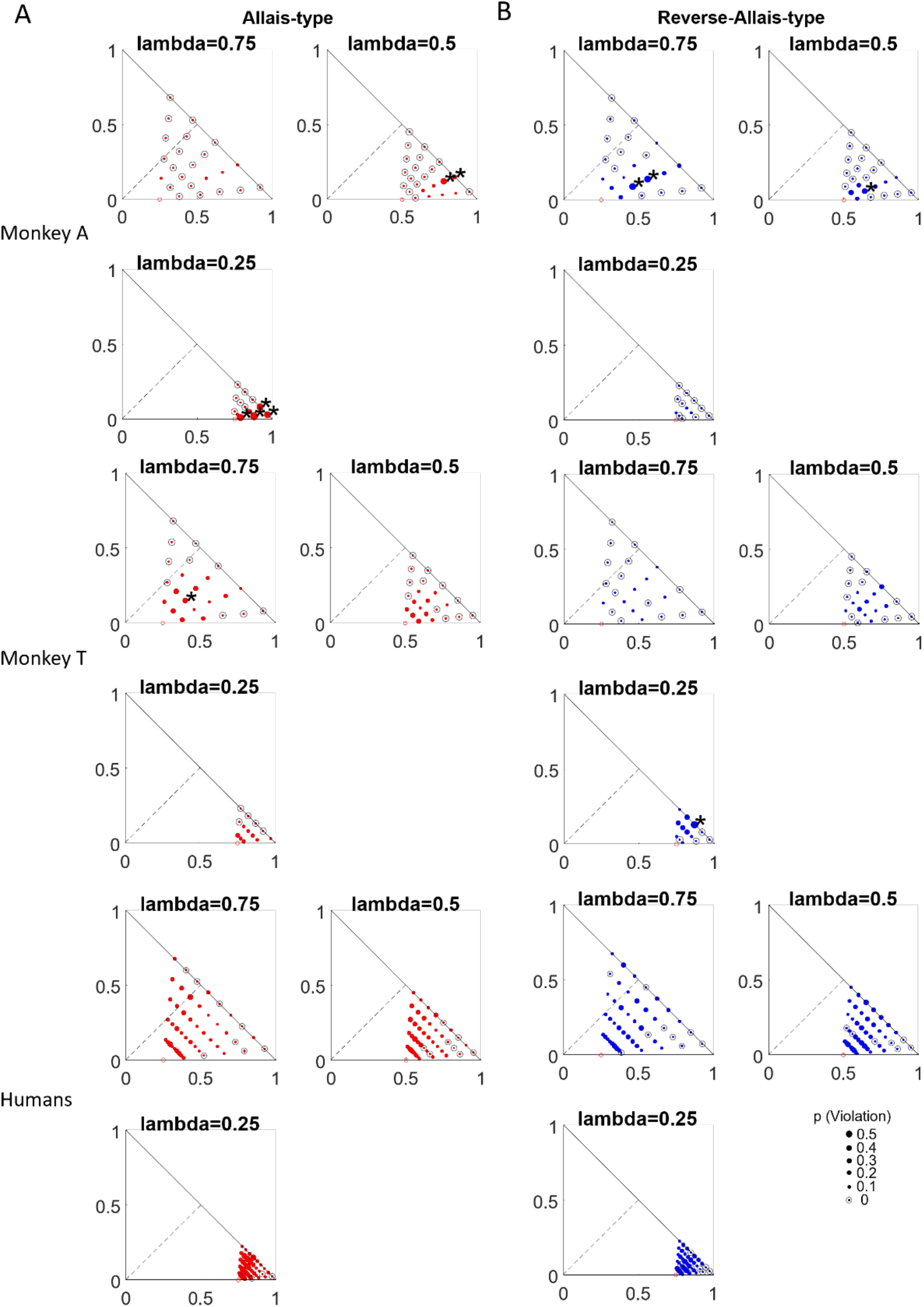
Violations of independence axiom across probabilities. Frequency and direction of the independence axiom violation across different probabilities. Dot size represents the probability of violation across (monkey) sessions or (human) participants, comparing different lambdas to the initial choice between options A and B (lambda = 1.0). (A) Red dots represent the probability of Allais-type violations. (B) Blue dots represent the probability of reverse-Allais-type violations. Hallow black circles indicate no violation for that choice set. * significance P < 0.05; binomial test in the two test sets. The correlation between predicted and actual violation was significant (GLM model with human data to predict monkey behavior, Spearman Rho = 0.201, P = 0.029), when lambda was used as the regressor.

**Figure S3.**
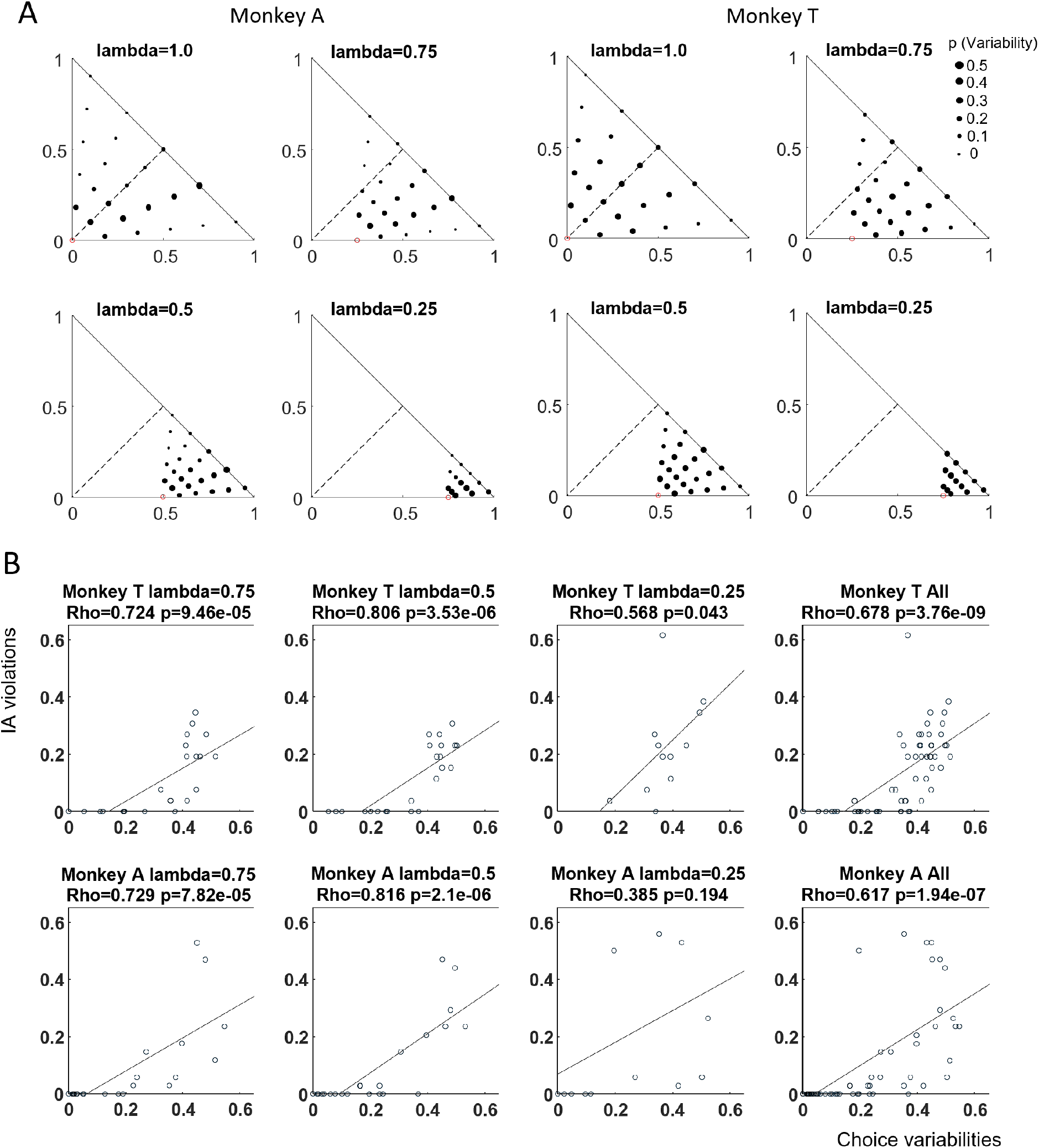
Choice variability partly explains independence axiom (IA) violations in monkeys. (A) Choice variability (standard deviation) across different probabilities. Size of dots representing the variability (standard deviation) of choices averaged across sessions (34 sessions for Monkey A and 26 sessions for Monkey T). (B) Correlation of choice variabilities (sum of two tests) and IA violations.

